# Efficiency and localisation of AURKA degradation by PROTACs is modulated by deubiquitinases UCHL5 and target-selective OTUD6A

**DOI:** 10.1101/2025.04.23.650020

**Authors:** Annabel Cardno, Karen Roberts, Catherine Lindon

**Author notes:** For correspondence: Annabel Cardno, Catherine Lindon.

## Abstract

Proteolysis-targeting chimeras (PROTACs) represent a promising new drug modality for novel therapeutics. However, the cellular mechanisms and regulatory pathways underlying their activity are not fully understood. Here, we unveil the role of deubiquitinases (DUBs) in regulating PROTAC activity, by screening 97 human DUBs for their influence on degradation of cell-cycle kinase AURKA using siRNA-mediated knockdown. Our findings reveal that DUBs OTUD6A and UCHL5 counteract degradation of AURKA by small molecule PROTACs. Further investigation using orthogonal dTAG PROTACs indicates that the PROTAC-opposing effect of OTUD6A is target-specific for AURKA, while UCHL5 counteracts degradation triggered by other PROTACs dependent on ubiquitin ligase adaptor CRBN, but not VHL. Furthermore, we show that differential sensitivity of the nuclear pool of AURKA to PROTAC-mediated degradation is fully explained by the specific subcellular localisation pattern of OTUD6A. These findings enhance our understanding of cellular pathways underpinning the action of PROTACs and indicate that combinations of DUB inhibitors and PROTACs will lead to enhanced target degradation and potential improvement in therapeutic outcomes.

## Introduction

Over recent years the field of targeted protein degradation (TPD) has exploded as a new therapeutic strategy, enabling selective degradation of proteins of interest (POIs). Proteolysis-targeting chimeras (PROTACs) are one of the leading modalities of TPD tools; PROTACs consist of distinct binding moieties for a POI and E3 ubiquitin ligase connected by a chemical linker. This induced proximity facilitates ubiquitination of the POI which serves as a signal for its proteasomal degradation^1^. The majority of PROTACs to date have been designed to recruit the cullin-RING ligases CRL4^CRBN^ and CRL2^VHL^, although expansion of the E3 landscape is a key priority for current research^2^. Over 20 PROTACs have now entered clinical trials, with advanced candidates estrogen receptor PROTAC ARV-471^3^ and androgen receptor degrader ARV-110^4^ under investigation in phase III clinical trials (NCT03888612, NCT05654623).

Despite the advancement of many PROTACs into clinical trials, the cellular mechanisms and regulatory pathways influencing PROTAC efficacy remain incompletely understood. Target-specific factors, including subcellular localisation of targets, have been shown to play a role in determining the amenability of certain substrates to PROTAC-mediated degradation^5^. For example, our previous work demonstrated the cell-cycle kinase AURKA shows differential sensitivity to PROTAC-mediated degradation depending on its subcellular localisation; AURKA on the mitotic spindle was degraded efficiently by CRBN-recruiting small molecule PROTAC (PROTAC-D) whilst AURKA at the centrosome was resistant to PROTAC-mediated degradation^6^. Other studies have also shown the importance of subcellular localisation of a target in determining its degradation by PROTACs^7,8^, although the mechanisms underpinning these observations remain largely unexplored.

Ubiquitination of substrates requires a coordinated cascade of ubiquitin-activating (E1), ubiquitin-conjugating (E2) and ubiquitin ligase (E3) enzymes^9^. It is widely accepted that K11/K48-linked ubiquitin chains serve as signals for proteasomal degradation^10,11^, although recent evidence highlighted the role of branched ubiquitin chains (e.g., K48/K63- and K29/K48-branched chains) in enhancing proteasomal degradation of substrates in response to PROTACs^12,13^. Beyond the ubiquitination machinery, other components of the ubiquitin proteasome system (UPS), such as the AAA+ ATPase p97^14^, are critical for ubiquitin chain processing and play a role in determining the fate of a PROTAC substrate^15^. The role of deubiquitinase enzymes (DUBs), in determining PROTAC efficacy remains largely unexplored. Given their ability to cleave ubiquitin chains from substrates and stabilise targets^16^, it is likely that DUBs could rescue PROTAC targets from proteasomal degradation.

In this study, we test the hypothesis that DUBs could oppose PROTAC-mediated degradation of cellular targets, using AURKA as a model target previously shown to exhibit differential PROTAC sensitivity dependent on subcellular localisation. We screen a panel of 97 DUBs by siRNA-mediated depletion to identify those which modulate PROTAC-sensitivity of AURKA. Our validation of screen results reveals DUBs with varying levels of specificity for both target protein and E3 and identifies one DUB, OTUD6A, which selectively protects the cytoplasmic pool of AURKA from PROTAC-mediated degradation.

## Results

### Forced nuclear localisation of AURKA enhances its sensitivity to degradation by PROTAC-D, independently of ternary complex formation

Previous studies from our lab demonstrated that distinct subcellular pools of AURKA exhibit differential sensitivity to degradation by PROTAC-D, an AURKA-targeting PROTAC that combines the AURKA inhibitor alisertib with pomalidomide^6^. Notably, we observed that an N-terminally truncated version of AURKA-Venus (Δ67) was simultaneously more efficiently degraded by PROTAC-D and more strongly localised to the nuclear compartment^6,17^.

To explore how nuclear localisation influences PROTAC-D-induced degradation of AURKA, we engineered an AURKA-Neon construct tagged with a nuclear localisation signal (NLS) and expressed it alongside wild-type (WT) AURKA-Neon in U2OS FZR1^KO^ cells (in which AURKA is insensitive to its cognate E3, the APC/C)^18^. Quantitative live-cell imaging confirmed the nuclear localisation of NLS-AURKA-Neon compared to WT-AURKA-Neon (**Figure 1A, B**).

**Figure 1:**
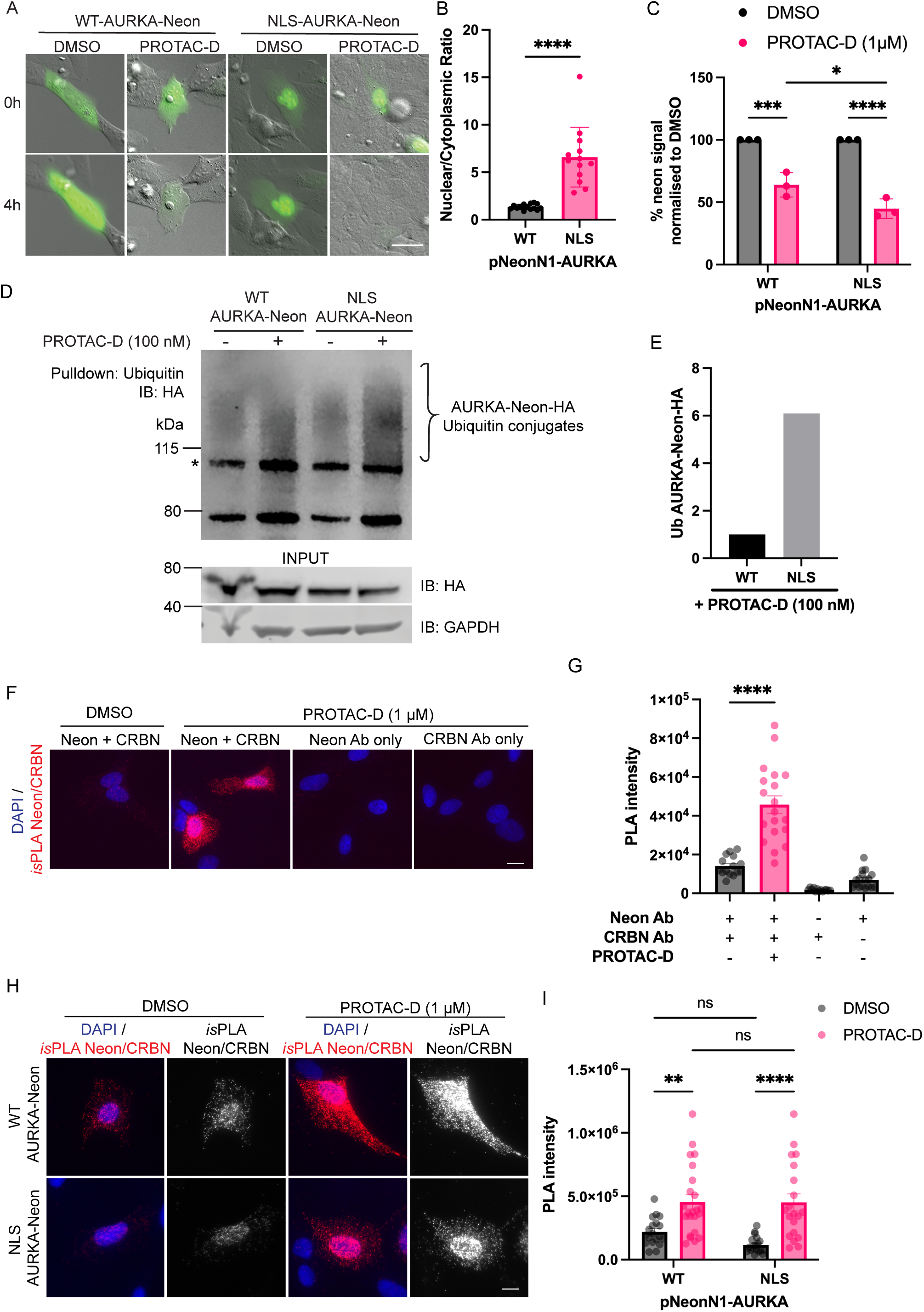
PROTAC-D treatment leads to greater degradation of nuclear AURKA compared to cytoplasmic AURKA. **A** Representative images of U2OS FZR1^KO^ cells transfected with pNeonN1-AURKA (wildtype, WT) or nuclear-localised pNeonN1-NLS-AURKA (NLS) and treated with DMSO or PROTAC-D (1 µM) for 4 hours. Images acquired by widefield fluorescence microscopy. Scale bar 10 µm. **B** Quantification of the nuclear-to-cytoplasmic fluorescence intensity ratio for WT- and NLS-AURKA-Neon constructs shown in (**A**). Data show pooled single-cell measurements with bars representing mean ± SD from one experiment (n ≥ 13 cells) and are representative of three biological repeats; statistical analysis by unpaired t-test. **C** DMSO-normalised mNeon fluorescence of WT- or NLS-AURKA-Neon in U2OS FZR1^KO^ cells after 4-hour treatment with DMSO or PROTAC-D (1 µM). Images were acquired using time-lapse microscopy and mNeon fluorescence levels in single cells measured at 0- and 4-hour timepoints. Substrate degradation is expressed as percentage change of fluorescence at 4 hours in single cells, normalised against the mean value from DMSO-treated cells and data points are mean values ± SD from three biological repeats (≥ 18 cells from multiple fields analysed per condition across all repeats). Statistical significance determined by two-way ANOVA with Tukey’s post hoc multiple comparison test. **D** U2OS FZR1^KO^ cells co-transfected with pNeonN1-AURKA/pNeonN1-NLS-AURKA and pcDNA3-FLAG-Ub were treated with DMSO or PROTAC-D (100 nM) for one hour. Ubiquitinated proteins from cell extracts were captured by pulldown with ubiquitin affinity beads. The band marked (*) is a non-specific band. **E** Quantification of PROTAC-D-induced ubiquitin conjugates shown in (**D**), expressed as the ratio of ubiquitin conjugates to input detected via the HA tag on mNeon, normalised to the ratio for WT-AURKA-Neon. The ubiquitin smear was quantified above the non-specific band at around 110 kDa, marked (*). **F** Representative images of mNeon/CRBN *is*PLA signal, indicative of ternary complex formation, detected in interphase U2OS FZR1^KO^ cells transfected with pNeonN1-AURKA. Cells were treated for 1 hour with DMSO or PROTAC-D (1 µM) and processed for *is*PLA using mNeon and CRBN primary antibodies in combination or alone to control for non-specific signal. Scale bar 10 µm. **G** Quantification of *is*PLA signal from (**F**). Total *is*PLA intensity was measured in single cells and pooled data plotted as mean ± SEM (n ≥ 13 cells). Statistical significance was assessed using one-way ANOVA with Tukey’s post hoc multiple comparisons test. **H** Representative images of mNeon/CRBN *is*PLA signal in interphase U2OS FZR1^KO^ cells transfected with pNeonN1-AURKA or pNeonN1-NLS-AURKA after 1 hour treatment with DMSO or PROTAC-D (1 µM). Scale bar 10 µm. **I** Quantification of *is*PLA signal from (**H**). Total *is*PLA intensity was measured in single cells and pooled data plotted as mean ± SEM (n ≥ 19 cells). Statistical significance was assessed using two-way ANOVA with Tukey’s post hoc multiple comparisons test.

Following PROTAC-D treatment, single-cell time-lapse imaging showed significantly greater degradation of NLS-AURKA-Neon than WT (**Figure 1A, C**). Ubiquitin pulldowns confirmed increased ubiquitination of NLS-AURKA-Neon, evidenced by an increase in the intensity of the ubiquitin smear, and upward shift of ubiquitinated NLS-AURKA-Neon to higher molecular weights (**Figure 1D, E, Supplementary Figure 1**).

PROTAC-D recruits the CRL4 adaptor protein CRBN. Given prior literature suggesting a stronger nuclear localisation of CRBN in certain cell types^19,20^, and correlations between ternary complex formation (TCF) and neosubstrate degradation^21,22^, we hypothesised that enhanced TCF in the nucleus might drive the increased degradation of nuclear-localised AURKA. To test this, we used the *in-situ* proximity ligation assay (*is*PLA) to measure TCF in single cells. U2OS FZR1^KO^ cells transfected with AURKA-Neon constructs were treated with DMSO or PROTAC-D, and *is*PLA performed to detect interactions between mNeon and endogenous CRBN. A robust PROTAC-D-dependent increase in single-cell *is*PLA signal was observed (**Figure 1F, G**). However, there was no difference in the *is*PLA signal measured when comparing WT- and NLS-AURKA-Neon transfected cells treated with PROTAC-D (**Figure 1H, I**), indicating that enhanced degradation of nuclear AURKA depends upon events downstream of TCF.

### Primary screen for DUBs modulating PROTAC-D activity

We concluded that differential degradation of subcellular pools of AURKA would depend upon post-ubiquitination regulation of the target, pointing to DUBs as candidate limiting factors in the activity of PROTAC-D. To identify DUBs influencing PROTAC-D activity, we conducted a systematic siRNA screen targeting 97 DUBs.

To facilitate high-throughput screening, we developed a doxycycline-inducible HiBiT-AURKA U2OS cell line (U2OS HiBiT-AURKA^TO^) for sensitive readout of AURKA levels in bioluminescence assays. The HiBiT-AURKA signal was dependent on treatment with doxycycline and addition of furimazine, the NanoLuc substrate, to the assay lytic buffer (**Figure 2A**). We confirmed that PROTAC-D and PROTAC-DX (a structural analogue of PROTAC-D differing only in linker-length) caused significant decrease in HiBiT-AURKA levels after 4 hours, which was rescued by the proteasome inhibitor MG132, confirming proteasome-dependent AURKA degradation by PROTACs in this assay (**Figure 2A**). The elevated levels of HiBiT-AURKA levels with MG132 compared to DMSO-treated cells likely reflects the normal proteasomal degradation of AURKA that occurs independently of PROTAC-D. The assay successfully reported on dose-dependent action of PROTAC-D and PROTAC-DX (**Figure 2B**). At concentrations above 1 µM, both PROTACs displayed the characteristic ‘hook effect’, where bioluminescence levels increased due to excessive binary binding of PROTACs to either AURKA or CRBN^23^. Importantly, no effect on cell viability was observed during the 4-hour treatment window, even at high doses of PROTAC (**Figure 2C**). Although PROTAC-DX showed greater degradation of AURKA (**Figure 2B**), PROTAC-D was chosen for the screen as it showed less degradation of AURKB at the four-hour timepoint (28% versus 60%) (**Supplementary Figure 2**).

**Figure 2:**
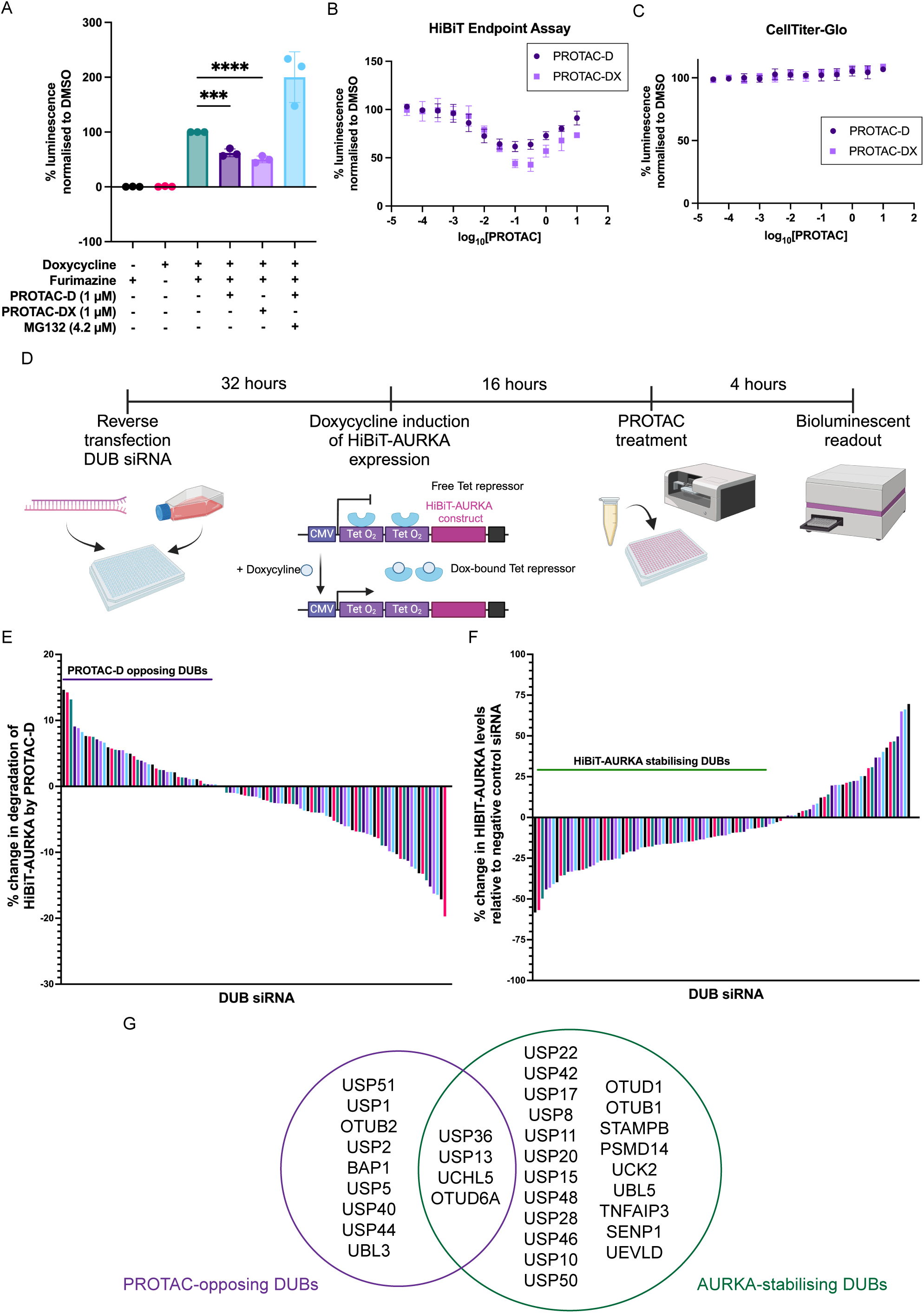
siRNA screen identifies DUBs regulating HiBiT-AURKA levels and sensitivity to PROTAC-D-mediated degradation. **A** HiBiT-AURKA bioluminescent signal normalised to DMSO controls in doxycycline-inducible U2OS HiBiT-AURKA^TO^ cells after 4-hour treatment with DMSO, PROTAC-D, PROTAC-DX or PROTAC-D plus proteasome inhibitor MG132, at the concentrations indicated. Individual data points represent mean values from three technical replicates per experiment, with bars showing mean value ± SD from three independent experiments (n=3). Statistical analysis by one-way ANOVA with Dunnett’s post-hoc multiple comparison test to DMSO. **B** Dose-response curve of DMSO-normalised HiBiT-AURKA bioluminescence, showing means ± SD from n=3 biological repeats. **C** Dose-response curve for the CellTiter-Glo®-based cell viability assay normalised to DMSO. Data represent means ± SD from n=3 biological repeats. **D** Workflow of the siRNA screen used to identify DUBs regulating HiBiT-AURKA levels. **E** Percentage change in PROTAC-D-mediated HiBiT-AURKA degradation over 4 hours, relative to negative control siRNA. Average values from two biological repeats are plotted. **F** Percentage change in HiBiT-AURKA levels in DMSO-treated cells relative to negative control siRNA. Average values from two biological repeats are plotted. **G** Summary of top-scoring DUB hits for those affecting PROTAC-D-mediated degradation (left) and those affecting basal HiBiT-AURKA levels (right).

Using a library of 97 siRNA pools, we individually depleted DUBs in U2OS HiBiT-AURKA^TO^ cells, then induced HiBiT-AURKA expression with doxycycline and treated the cells with DMSO or PROTAC-D for 4 hours before taking bioluminescence measurements (**Figure 2D**). Comparing PROTAC-induced degradation under DUB siRNA-treated and negative control siRNA-treated conditions revealed the DUBs affecting sensitivity of AURKA to PROTAC-D (**Figure 2E**). Unexpectedly, a large number of DUB siRNAs caused a decrease in basal HiBiT-AURKA levels in DMSO-treated cells, such that sensitivity to PROTAC-D appeared reduced in our assay (**Figure 2F**). This is consistent with the high turnover of AURKA in cells, suggesting that these DUBs may either directly remove ubiquitin from AURKA under basal conditions to protect AURKA from proteasomal degradation, or have roles in cellular pathways that indirectly affect AURKA levels. A Venn-diagram summarises the top-scoring hits (those affecting AURKA levels ≥3-fold SD of negative control) across these conditions (**Figure 2G**).

### Identifying DUBs which regulate endogenous AURKA expression and degradation by PROTAC-D

To further validate the role of the identified DUBs in regulating PROTAC-D activity, we selected a small number of candidate DUBs for further validation and ranked them according to their influence on PROTAC-D sensitivity (**Figure 3A**).

**Figure 3:**
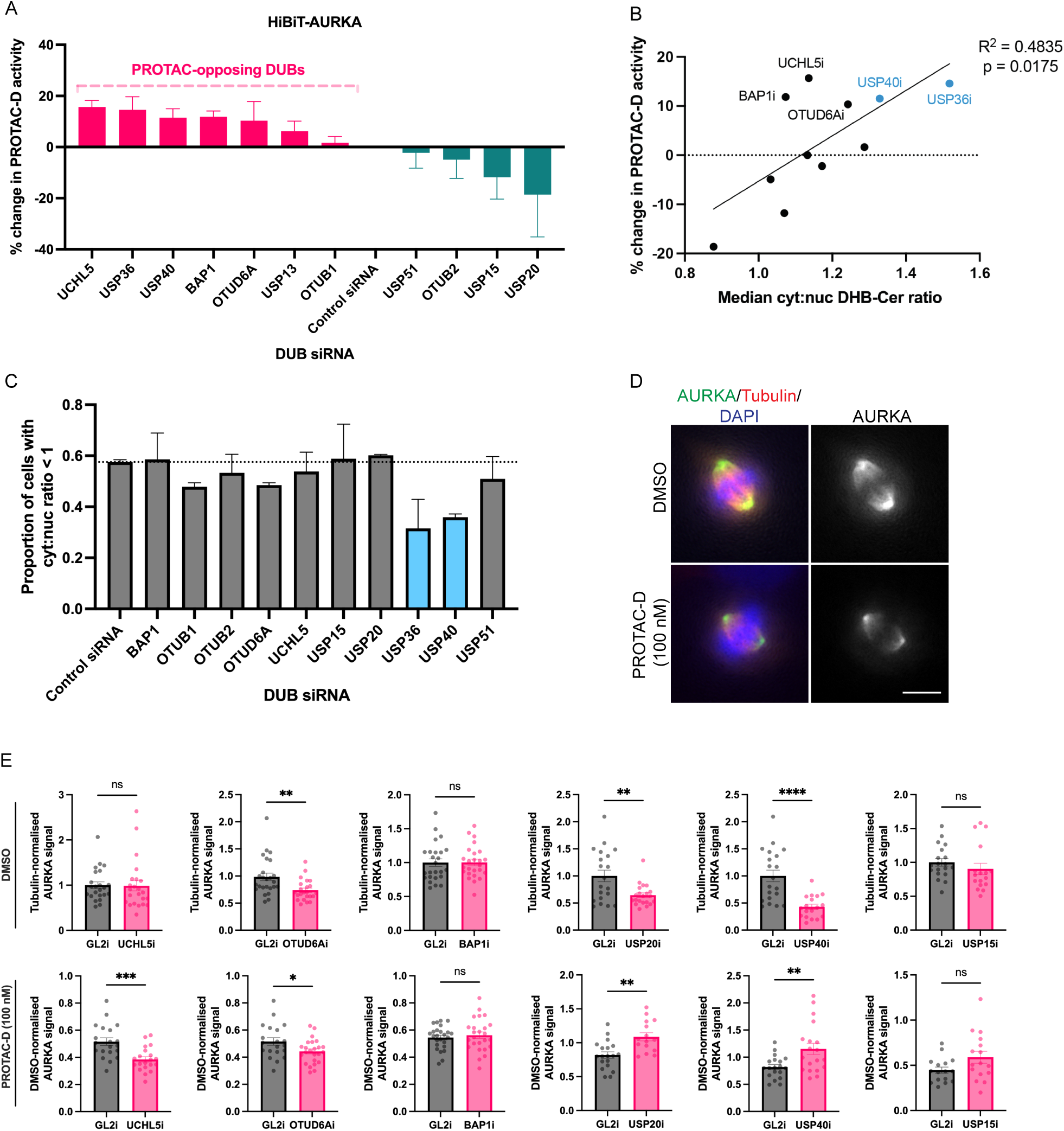
validation of DUB hits from siRNA screen as PROTAC-opposing DUBs. **A** U2OS HiBiT-AURKA^TO^ cells transfected with siRNA pools (4 oligos) targeting the indicated DUBs were treated with DMSO or PROTAC-D (100 nM) for 4 hours. HiBiT-AURKA bioluminescence was quantified, and PROTAC-D-mediated degradation was normalised to DMSO controls. Data are plotted as the percent change in degradation relative to negative control siRNA, with means ± SD from n=3 biological repeats. **B, C** U2OS^CDK2^ cells were transfected with siRNA pools targeting the indicated DUBs. 48 hours post-transfection nuclear-to-cytoplasmic DHB-Cer signal ratios were measured from widefield fluorescence images of live cells. **B** Data shows median DHB-Cer cytoplasmic:nuclear ratio plotted against percentage change in PROTAC-D activity as determined in (**A**), for each candidate DUB siRNA. Medians are the average values from two biological repeats, percentage changes in PROTAC-D activity are the average values from three biological repeats. Simple linear regression is plotted with Pearson’s statistical test for correlation. **C** Bars show the proportion of cells in each DUB siRNA-transfected pool with a cytoplasmic:nuclear ratio < 1, plotted as mean values ± SD from n=2 biological repeats, with ≥36 cells analysed per experiment. **D** Representative images of U2OS cells transfected with negative control siRNA and treated with DMSO or PROTAC-D (100 nM), stained for AURKA (green) and tubulin (red). DNA stained with DAPI. Scale bar 10 µm. **E** Quantification of whole-cell AURKA levels in pre-anaphase mitotic U2OS cells such as the examples shown in (**D**), transfected for 48 hours with indicated siRNA and treated with DMSO (upper panels) or PROTAC-D (lower panels) for 4 hours. AURKA intensity was measured in single cells and normalised to tubulin intensity. Data clouds show values from individual cells, with bars indicating mean values ± SEM for ≥ 15 cells analysed per condition. Statistical significance assessed via t-test.

Given that AURKA activity and stability fluctuates through the cell cycle according to presence of activating binding partners (highest in G2 and M phases) and activity of APC/C^FZR1^, which is higher in G1 phase than G2^24^, we sought to exclude DUBs whose knockdown might affect PROTAC-D activity in our screen indirectly due to changes in cell cycle progression. We employed U2OS^CDK2^ cells, a stable cell line expressing a cerulean-tagged CDK2 activity sensor, which allows visualisation of cell cycle phase based on localization of DHB-Cer^25^ (**Supplementary Figure 3A**).

Correlation analysis between the median cytoplasmic:nuclear DHB-Cer ratio (which increases with cell cycle progression) and changes in PROTAC-D sensitivity of HiBiT-AURKA revealed a significant positive correlation (p=0.0175) (**Figure 3B**). This supported our hypothesis that DUB knockdowns affecting cell cycle progression would confound measurements of PROTAC-D sensitivity. For example, USP40 and USP36 knockdowns led to both increased cytoplasmic:nuclear DHB-Cer ratios, indicative of shortened G1 phase, (**Figure 3C**) and increased PROTAC-D activity against AURKA (**Figure 3A**). This correlation was consistent with the idea that higher levels of a target protein show greater sensitivity to PROTACs^26,27^ but could also reflect increasing AURKA activity at later stages of the cell cycle, with associated increase in accessibility of the ATP-binding pocket to PROTAC binding. We tested the effect of cell cycle progression on PROTAC-D activity by synchronising cell populations before PROTAC treatment and found a small but not significant increase in PROTAC-D-mediated degradation of AURKA in G2- or M-phase-synchronised cells, while serum-starved cells (G0/G1-enriched) showed a significant reduction in degradation (**Supplementary Figure 3B**). Given this result, together with the marked effect of its knockdown on cytoplasmic:nuclear DHB-Cer ratio consistent with G2 phase arrest (**Figure 3B**), we excluded USP36 from subsequent studies. Since USP40 knockdown had a less dramatic effect on cytoplasmic:nuclear DHB-Cer ratio (**Figure 3B**), and is a little-known DUB, we included it in further analyses.

We note that a recent survey of DUB activities has identified USP36 as a ’high impact’ DUB in targeting almost 20% of the tested pool of ubiquitinated proteins, it may therefore prove an interesting candidate for future investigation of PROTAC-opposing DUBs^28^.

We assessed the impact of our remaining DUB hits on endogenous AURKA levels through immunofluorescence analysis of pre-anaphase mitotic cells, where AURKA is most biologically active (**Figure 3D**). We found that knockdown of OTUD6A, USP20 or USP40 led to a reduction in AURKA levels even in the absence of PROTAC treatment (**Figure 3E**), pointing to these DUBs as potential regulators of mitotic AURKA. OTUD6A has previously been identified as a DUB that deubiquitinates AURKA^29^ and USP20 was reported as a hit from a previous screen for DUBs regulating AURKA^30^. To our knowledge, USP40 has not previously been shown to regulate AURKA. In presence of PROTAC-D treatment, we found reduced AURKA signal in the majority of experiments compared to the DMSO control, with significantly enhanced loss of endogenous AURKA after knockdown of OTUD6A or UCHL5, pointing to a potential role for these DUBs in removing PROTAC-D-generated ubiquitin chains from AURKA. Knockdown of BAP1 and USP15 had no effect on endogenous AURKA levels, despite their effects on exogenously expressed protein (**Figure 3A, B**), whereas USP20 and USP40 siRNA unexpectedly appeared to reduce activity of PROTAC-D against endogenous AURKA (**Figure 3E**). Overall, UCHL5 and OTUD6A emerged as the strongest candidate PROTAC-D-opposing activities, making them promising DUBs for further investigation.

### UCHL5 opposes PROTAC-mediated degradation of AURKA and other CRBN-targeting substrates

To further characterise the role of UCHL5 in counteracting PROTAC activity, we evaluated the effect of UCHL5 knockdown on degradation of AURKA by other commercially available AURKA PROTACs. Specifically, we tested JB170, an AURKA-specific PROTAC comprised of alisertib linked to thalidomide^31^, and TL12-186, a CRBN-dependent multi-kinase degrader^32^. TL12-186 showed the most potent degradation of AURKA at 100 nM, while JB170 caused the least degradation (**Figure 4A**). We tested the knockdown of UCHL5 using two individual oligos (**Figure 4B**) and proceeded with oligo #2 which demonstrated superior knockdown efficiency. UCHL5 siRNA treatment led to enhanced degradation of HiBiT-AURKA in the presence of each of the three PROTACs (**Figure 4C**). To rule out potential off-target siRNA effects, we engineered UCHL5-Neon constructs with siRNA-resistant sequences and transfected them alongside UCHL5 siRNA into U2OS HiBiT-AURKA^TO^ cells. The enhancement of PROTAC-D and TL12-16 activity upon UCHL5 knockdown was fully rescued by expression of siRNA-resistant UCHL5, confirming the specificity of UCHL5 siRNA (**Figure 4D**). The best characterised function of UCHL5 is its association with the proteasome^33^ and our results are consistent with the hypothesis that UCHL5 removes ubiquitin chains from substrates recruited to the proteasome, hence limiting their degradation (and in our experiments, the activity of AURKA PROTACs). Overexpression of UCHL5 did not affect PROTAC-D activity in our experiments (**Supplementary Figure 4A**), consistent with the idea that the PROTAC-relevant pool of UCHL5 is stoichiometrically associated with the proteasome^33,34^.

**Figure 4:**
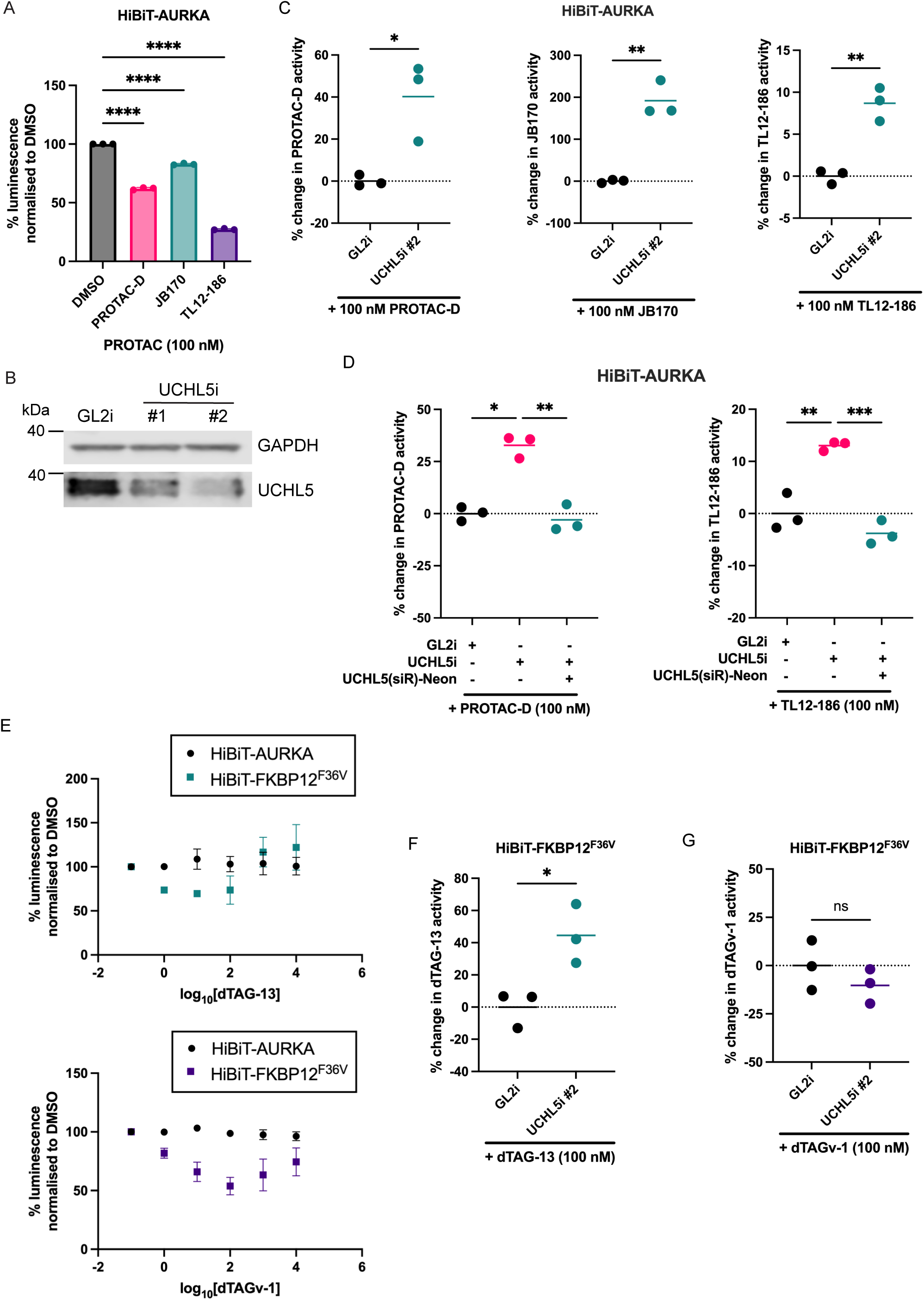
UCHL5 opposes PROTAC-mediated degradation of AURKA and other substrates. **A** DMSO-normalised HiBiT-AURKA levels after 4-hour treatments with DMSO, PROTAC-D, JB170, or TL12-186 (100 nM) in U2OS HiBiT-AURKA^TO^ cells. Data represent means ± SD from n=3 biological repeats; statistical analysis by one-way ANOVA with Dunnett’s post-hoc multiple comparisons test to DMSO-treated cells. **B** Immunoblot analysis of UCHL5 levels in U2OS HiBiT-AURKA^TO^ cells transfected with negative control siRNA (GL2i) or UCHL5-targeting siRNAs (two oligos). Lysates were collected 48 hours post-transfection. **C** U2OS HiBiT-AURKA^TO^ cells transfected with GL2 siRNA or UCHL5 siRNA were treated with DMSO, PROTAC-D, JB170 or TL12-186 (100 nM). HiBiT-AURKA bioluminescence signal was normalised to DMSO-treated controls for each transfection condition. Percentage change in PROTAC-mediated HiBiT-AURKA degradation relative to negative control siRNA is plotted, with individual data points corresponding to mean values of technical replicates from n=3 biological repeats and line to indicate the mean value of biological repeats; statistical analysis by unpaired t-test. **D** siRNA-rescue experiment in U2OS HiBiT-AURKA^TO^ cells transfected with combinations of GL2 siRNA, UCHL5 siRNA, and siRNA-resistant pNeonN1-UCHL5 (siR). Cells were treated with DMSO, PROTAC-D, or TL12-186 (100 nM) for 4 hours, and HiBiT-AURKA signal was normalised to DMSO-treated controls. Percentage change in PROTAC-mediated HiBiT-AURKA degradation relative to negative control siRNA is plotted. Data points correspond to mean values of technical replicates from n=3 biological repeats with line to indicate the mean value of biological repeats; statistical analysis by one-way ANOVA with Tukey’s post hoc multiple comparisons test. **E** Dose-response curves of U2OS cells transfected with pNeonN1-HiBiT-FKBP12^F36V^ or pcDNA3.1-HiBiT-AURKA and treated with varying doses of dTAG-13 (upper panel) or dTAGv-1 (lower panel). Data are DMSO-normalised means ± SD from n=3 biological repeats. **F, G** U2OS cells transfected with pNeonN1-HiBiT-FKBP12^F36V^ and GL2 siRNA or UCHL5 siRNA were treated with DMSO, dTAG-13 **(F)**, or dTAGv-1 **(G)** at the indicated concentrations for 4 hours. HiBiT bioluminescent signal was normalised to DMSO-treated controls. Percentage change in PROTAC-mediated HiBiT-FKBP12^F36V^ degradation in UCHL5 siRNA relative to negative control siRNA is plotted. Data points correspond to mean values of technical replicates from n=3 biological repeats with line to indicate the mean value of biological repeats; statistical analysis by unpaired t-test.

Next, we investigated whether the PROTAC-opposing effect of UCHL5 was selective for AURKA, or for CRBN-recruiting PROTACs, or for neither. To test this, we utilised the dTAG system^35,36^, whereby orthogonal dTAG PROTACs target FKBP12^F36V^-tagged constructs for degradation via either CRBN (dTAG-13) or VHL (dTAGv-1). HiBiT-FKBP12^F36V^ constructs were generated and validated, showing dose-dependent degradation that was dependent on the FKBP12^F36V^ tag. The hook effect was seen at higher concentrations (**Figure 4E**). Knockdown of UCHL5 enhanced the degradation of HiBiT-FKBP12^F36V^ by dTAG-13, indicating that UCHL5’s PROTAC-opposing effect extends beyond AURKA-targeting PROTACs to other CRBN-recruiting systems (**Figure 4F**). However, knockdown of UCHL5 had no impact on the degradation of HiBiT-FKBP12^F36V^ by dTAGv-1, a VHL-recruiting PROTAC (**Figure 4G**). The differential effect of UCHL5 on CRBN-versus VHL-mediated dTAG-induced degradation may be attributed to variations in extent and topology of ubiquitin chains conferred by the distinct E3s that would affect the rate of processing of substrate at the proteasome. We hypothesised that VHL might confer more extensive (higher MW) ubiquitin chains but did not find any evidence for this in a ubiquitin pulldown experiment (**Supplementary Figure 4B, C**).

### OTUD6A opposes PROTAC-mediated degradation of AURKA

Next, we investigated the role of OTUD6A in counteracting PROTAC activity. Knockdown of OTUD6A was validated using two independent siRNA oligos (**Figure 5A**) and shown to enhance the degradation of AURKA by both PROTAC-D and TL12-186 (**Figure 5B**). The results for JB170 were less consistent, as only one siRNA oligo produced a PROTAC-enhancing effect (**Figure 5B**). To confirm the specificity of the OTUD6A siRNA, we engineered an siRNA-resistant OTUD6A-Neon construct. Co-transfection of U2OS HiBiT-AURKA^TO^ with this construct and OTUD6A siRNA fully reversed the PROTAC-enhancing effects observed with OTUD6A knockdown (**Figure 5C**), confirming the specificity of OTUD6A siRNA.

**Figure 5:**
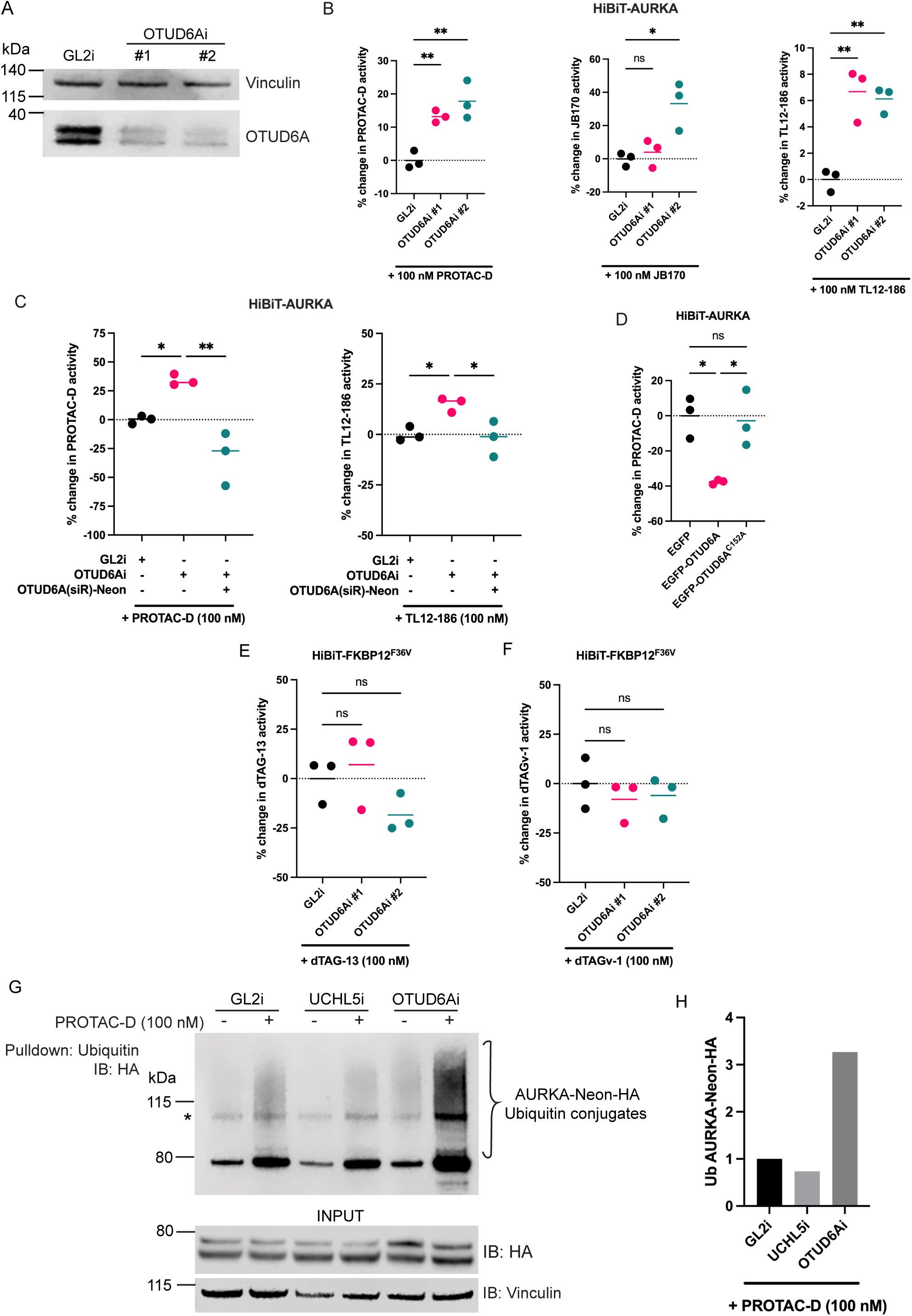
OTUD6A opposes PROTAC-mediated degradation of AURKA in a target-specific manner. **A** Immunoblot analysis of U2OS HiBiT-AURKA^TO^ cells transfected with negative control siRNA (GL2i) or OTUD6A-targeting siRNAs (two oligos). Lysates were collected 48 hours post-transfection. **B** U2OS HiBiT-AURKA^TO^ cells transfected with GL2 siRNA or OTUD6A siRNA were treated with DMSO, PROTAC-D, JB170 or TL12-186 (100 nM). HiBiT-AURKA signal was normalised to DMSO-treated controls for each transfection condition. Data points show percentage change in PROTAC-mediated HiBiT-AURKA degradation relative to negative control siRNA and correspond to mean values of technical replicates from n=3 biological repeats, with line to indicate the mean value of biological repeats; statistical analysis by one-way ANOVA with Tukey’s post hoc multiple comparisons test. **C** siRNA-rescue experiment in U2OS HiBiT-AURKA^TO^ cells transfected with combinations of GL2 siRNA, OTUD6A siRNA, and siRNA-resistant pNeonN1-OTUD6A (siR). Cells were treated with DMSO, PROTAC-D, or TL12-186 (100 nM) for 4 hours, and HiBiT-AURKA signal was normalised to DMSO-treated controls. Data points show percentage change in PROTAC-mediated HiBiT-AURKA degradation relative to negative control siRNA and correspond to mean values of technical replicates from n=3 biological repeats, with line to indicate the mean value of biological repeats; statistical analysis by one-way ANOVA with Tukey’s post hoc multiple comparisons test. **D** U2OS HiBiT-AURKA^TO^ cells transfected with pEGFP, pEGFP-OTUD6A, or catalytically inactive pEGFP-OTUD6A^C152A^ plasmids were treated with DMSO or PROTAC-D (100 nM) for 4 hours and HiBiT-AURKA signal was normalised to DMSO-treated condition. Data points show percentage change in PROTAC-mediated HiBiT-AURKA degradation relative to pEGFP overexpression and correspond to mean values of technical replicates from n=3 biological repeats, with line to indicate the mean value of biological repeats; statistical analysis by one-way ANOVA with Tukey’s post hoc multiple comparisons test. **E, F** U2OS cells transfected with pNeonN1-HiBiT-FKBP12^F36V^ and GL2 or OTUD6A siRNA were treated with DMSO, dTAG-13 (**E**), or dTAGv-1 (**F**) at the indicated concentrations for 4 hours. Data points show percentage change in PROTAC-mediated HiBiT-FKBP12^F36V^ degradation relative to negative control siRNA, plotted as mean values of technical replicates from n=3 biological repeats, with line to indicate the mean value of biological repeats; statistical analysis by unpaired t-test. **G** U2OS FZR1^KO^ cells co-transfected with pNeonN1-AURKA-Neon-HA, pcDNA3-FLAG-Ub and GL2, OTUD6A or UCHL5 siRNA were treated with DMSO or PROTAC-D (100 nM) for one hour. Ubiquitinated proteins from cell extracts were captured by pulldown with ubiquitin affinity beads. **H** Quantification of (**G**): chart shows the ratio of ubiquitin conjugates to input after PROTAC-D treatment for each condition, normalised to the ratio for GL2i. The quantified ubiquitin smear region was above the non-specific band at around 110 kDa, marked (*).

Overexpression of OTUD6A caused a significant decrease in PROTAC activity (**Figure 5D**), an effect abolished by mutation of the catalytic site (OTUD6A^C152A^). We concluded that OTUD6A acts to stabilise AURKA against PROTAC-mediated degradation through its deubiquitination activity.

We next assessed, using dTAG PROTACs, whether OTUD6A’s PROTAC-opposing effect was specific to AURKA-targeting PROTACs or to CRBN-harnessing tools, or to neither. In contrast to UCHL5, OTUD6A knockdown had no significant effect on HiBiT-FKBP12^F36V^ degradation mediated by either CRBN-recruiting dTAG-13 (**Figure 5E**) or VHL-recruiting dTAGv-1 (**Figure 5F**). We concluded that OTUD6A’s impact on PROTAC activity is specific to AURKA-targeting PROTACs.

In summary, while UCHL5 appears to exhibit generalised PROTAC-opposing activity against targets of CRBN-harnessing PROTACs, OTUD6A’s effect appears to selectively regulate AURKA degradation by PROTACs. We tested the effect of the two DUBs in ubiquitin pulldowns and found a striking enhancement of PROTAC-induced ubiquitin conjugates associated with AURKA-Neon-HA in OTUD6A siRNA conditions, suggesting OTUD6A removes ubiquitin from AURKA, thereby protecting it from proteasomal degradation (**Figure 5G, H**). UCHL5 siRNA had no such effect, raising questions about the precise mechanism through which it exerts its PROTAC-protective effect.

### Enhanced nuclear degradation of AURKA in response to PROTACs is target-specific

Having identified OTUD6A as an AURKA-specific PROTAC-opposing DUB, we set out to investigate whether it might contribute to compartment-specific modulation of PROTAC activity. First, we examined whether the enhanced degradation of nuclear-localised AURKA upon PROTAC-D treatment (shown in Figure 1) was a property of PROTAC-D or of AURKA-targeting PROTACs, or a generalised feature of PROTAC activity. We engineered a series of constructs containing AURKA, FKBP12^F36V^, mNeon and NLS sequences, targetable by dTAG PROTACs (**Figure 6A**). The expression of these artificial neosubstrates and their FKBP12^F36V^-dependent degradation in presence of dTAG-13 were quantified by live-cell imaging (**Figure 6B, 6C**).

**Figure 6:**
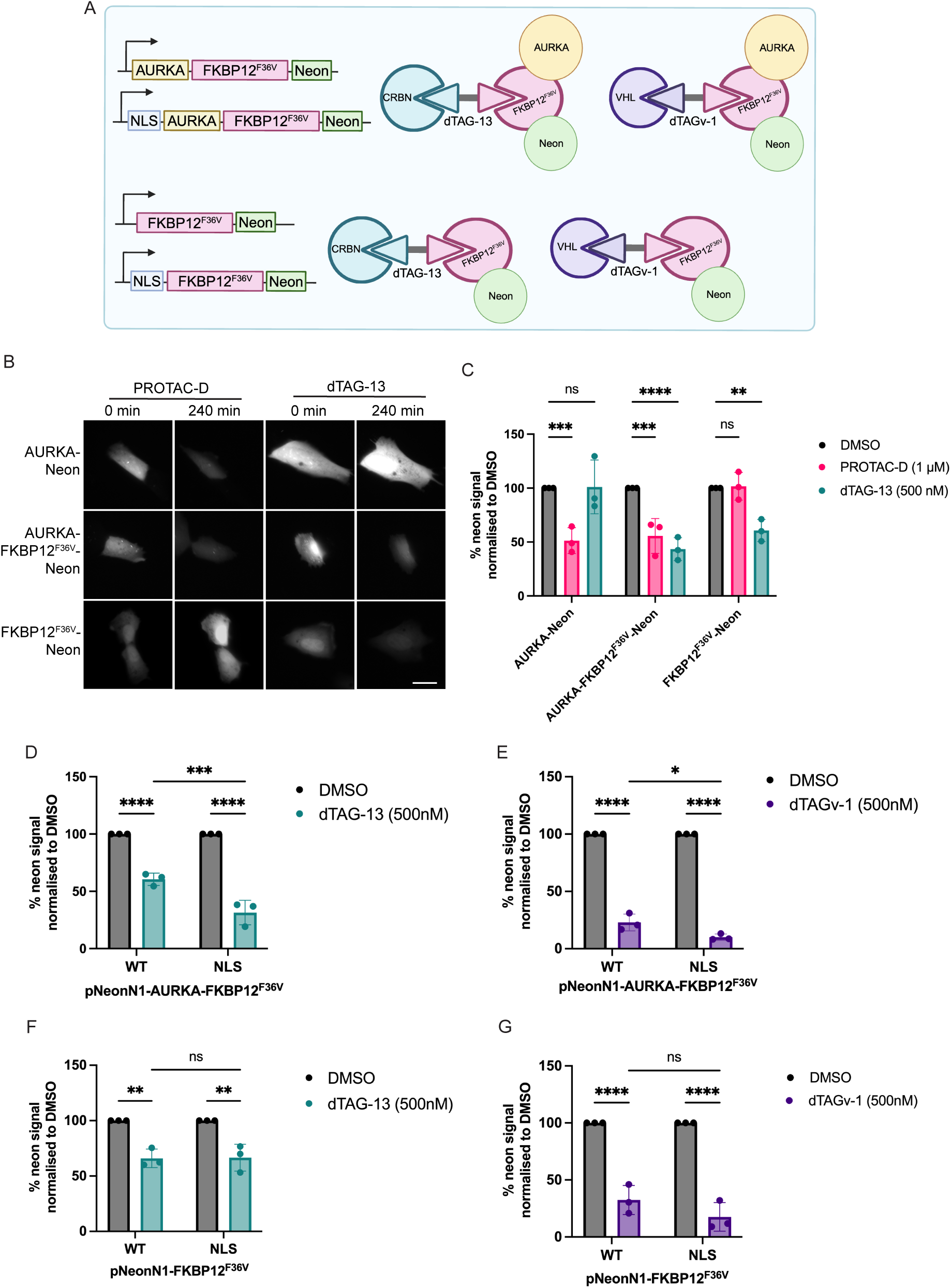
Enhanced degradation of the nuclear pool of AURKA in response to PROTACs is substrate-specific. **A** Schematic of FKBP12^F36V^-Neon constructs and engagement with CRBN-recruiting dTAG-13 and VHL-recruiting dTAGv-1 PROTACs. **B** Representative fluorescence images of U2OS FZR1^KO^ cells transfected with pNeonN1-AURKA, pNeonN1-AURKA-FKBP12^F36V^ or pNeonN1-FKBP12^F36V^, treated with PROTAC-D (1 µM) or dTAG-13 (500 nM) for 4 hours. Scale bar 10 µm. **C** Quantification of experiments illustrated in (**B**). Images were acquired using time-lapse microscopy and mNeon fluorescence levels in single cells measured at 0- and 4-hour timepoints. Substrate degradation is expressed as percentage change of fluorescence at 4 hours in single cells, normalised against the mean value from DMSO-treated cells. Bar chart shows mean values ± SD from three biological repeats, with ≥ 15 cells from multiple fields analysed per condition across all repeats. Statistical significance determined by two-way ANOVA with Tukey’s post hoc multiple comparison test. **D, E** U2OS FZR1^KO^ cells were transfected with WT- or NLS-AURKA-FKBP12^F36V^-Neon and subjected to timelapse fluorescence imaging after treatment with DMSO, dTAG-13 (**D**) or dTAGv-1 (**E**) at the indicated concentrations. mNeon fluorescence was quantified in single cells at 0- and 4-hour timepoints and change in fluorescence at 4 hours normalised for the mean values from DMSO-treated cells. Bar chart shows mean values ± SD from three biological repeats, with ≥ 20 cells from multiple fields analysed per condition across all repeats. Statistical significance determined by two-way ANOVA with Tukey’s post hoc multiple comparison test. **F, G** U2OS FZR1^KO^ cells were transfected with WT- or NLS-FKBP12^F36V^-Neon and subjected to timelapse fluorescence imaging after treatment with DMSO, dTAG-13 (**F**) or dTAGv-1 (**G**) at the indicated concentrations. mNeon fluorescence was quantified in single cells at 0- and 4-hour timepoints and change in fluorescence at 4 hours normalised for the mean values from DMSO-treated cells. Bar chart shows mean values ± SD from three biological repeats, with ≥ 22 cells from multiple fields analysed per condition across all repeats. Statistical significance determined by two-way ANOVA with Tukey’s post hoc multiple comparison test.

We compared the degradation of WT-AURKA-FKBP12^F36V^-Neon and NLS-AURKA-FKBP12^F36V^-Neon over a 4-hour period. NLS-AURKA-FKBP12^F36V^-Neon was degraded more efficiently than WT-AURKA-FKBP12^F36V^-Neon by either dTAG-13 (**Figure 6D**) or dTAGv-1 (**Figure 6E**). We concluded that the enhanced degradation of nuclear AURKA described in **Figure 1** was not due to increased binding of PROTAC-D to the nuclear pool of AURKA (since the same effect was seen with a different target warhead) and was independent of the E3 ligase harnessed (since it was observed with both CRBN- and VHL-recruiting PROTACs).

Next, we compared the degradation of WT-FKBP12^F36V^-Neon and NLS-FKBP12^F36V^-Neon constructs lacking AURKA. These constructs were degraded to a similar extent by dTAG-13 (**Figure 6F**) and dTAGv-1 (**Figure 6G**). We therefore concluded that the enhanced nuclear degradation of AURKA by PROTACs is target-specific.

### Enhanced degradation of nuclear AURKA is dependent on OTUD6A

Upon overexpression of GFP-OTUD6A in U2OS cells, we observed that localisation of OTUD6A was predominantly cytoplasmic, as previously described (**Figure 7A**)^37^. We hypothesised that OTUD6A might act to selectively protect the cytoplasmic pool of AURKA from PROTAC-mediated degradation.

**Figure 7:**
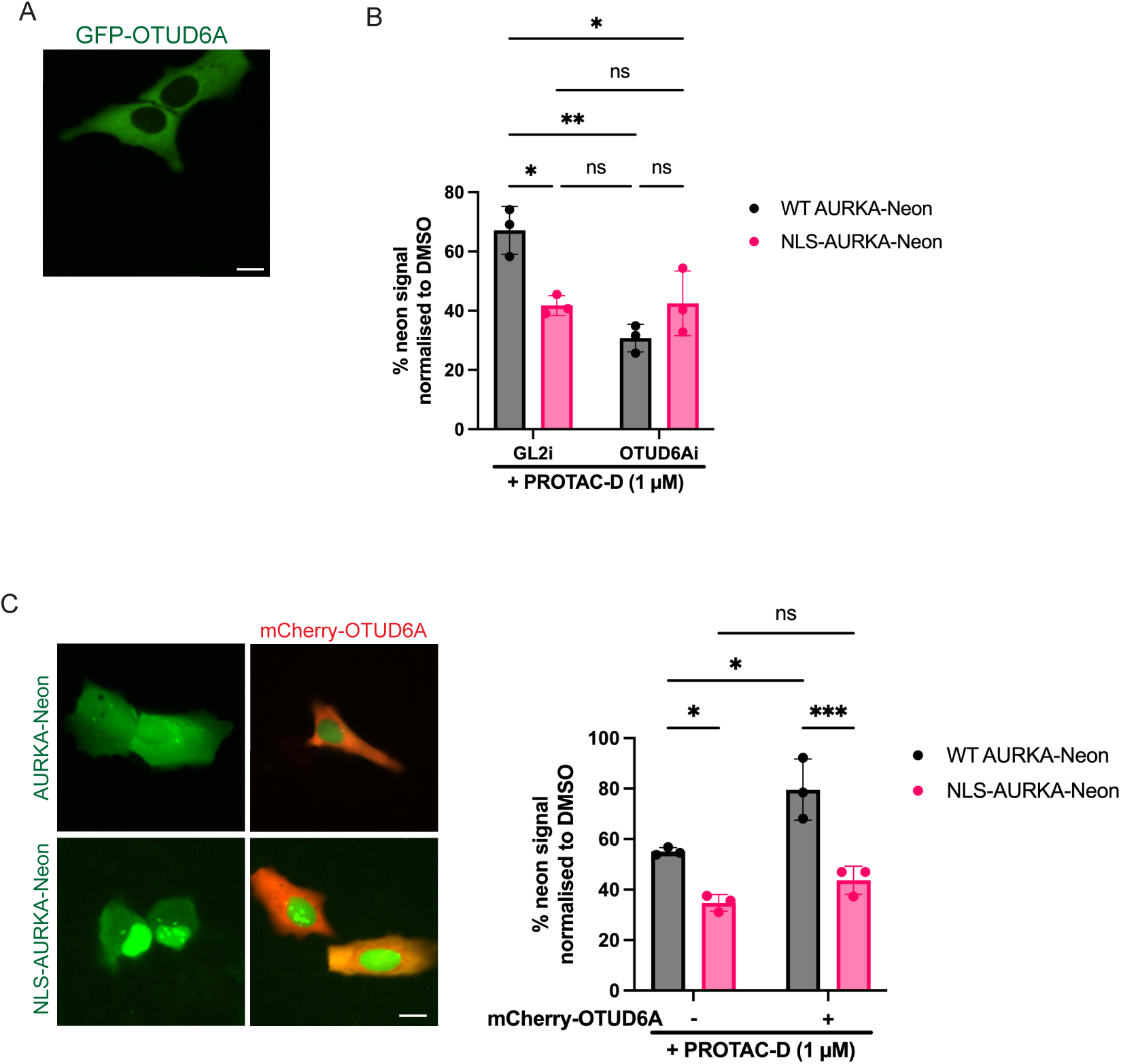
Enhanced degradation of the nuclear pool of AURKA is OTUD6A-dependent. **A** U2OS cells transfected with pEGFP-OTUD6A, showing subcellular localisation of GFP-OTUD6A in interphase cells. Scale bar 10 µm. **B** U2OS FZR1^KO^ cells were transfected with WT- or NLS-AURKA-Neon and GL2 siRNA or OTUD6A siRNA for 48 hours then treated for 4 hours with DMSO or PROTAC-D (1 µM) and subjected to timelapse fluorescence imaging. mNeon fluorescence was quantified in single cells at 0- and 4-hours and change in fluorescence normalised to the mean value from DMSO-treated cells. Bar charts show mean values ± SD from three biological repeats, with ≥ 17 cells from multiple fields analysed per condition across all repeats. Statistical significance determined by two-way ANOVA with Tukey’s post hoc multiple comparison test. **C** U2OS FZR1^KO^ cells were transfected with WT- or NLS-AURKA-Neon, with or without pN1-mCherry-OTUD6A for 24 hours then treated for 4 hours with DMSO or PROTAC-D (1 µM) and subjected to timelapse fluorescence imaging. mNeon fluorescence was quantified in single cells at 0- and 4-hours and change in fluorescence normalised to the mean value from DMSO-treated cells. Image panels show transfected cells at timepoint 0. Bar chart shows mean values ± SD from three biological repeats, with ≥ 14 cells from multiple fields analysed per condition across all repeats. Statistical significance determined by two-way ANOVA with Tukey’s post hoc multiple comparison test.

To test our hypothesis, we measured the degradation of WT-AURKA-Neon and NLS-AURKA-Neon by PROTAC-D in U2OS FZR1^KO^ cells under conditions of OTUD6A knockdown or overexpression. We found that OTUD6A knockdown significantly enhanced degradation of WT-AURKA-Neon, whereas degradation of NLS-AURKA-Neon was not affected (**Figure 7B**). Moreover, under OTUD6A siRNA conditions the enhanced degradation of NLS-AURKA-Neon compared to WT-AURKA-Neon was abolished.

Under conditions of OTUD6A overexpression there was significantly reduced degradation of WT-AURKA-Neon by PROTAC-D but, again, there was no effect on degradation of NLS-AURKA-Neon (**Figure 7C**).

From these results we conclude that OTUD6A protects the cytoplasmic pool of AURKA from PROTAC-mediated degradation and that nuclear-localised AURKA is insensitive to OTUD6A levels. Indeed, our experiments show that the presence of OTUD6A is sufficient to explain enhanced degradation of the nuclear pool of wild-type AURKA. Our results therefore demonstrate that the subcellular compartmentalisation of ubiquitin pathway components, such as OTUD6A, can drive differential degradation of PROTAC targets depending on their subcellular localisation.

## Discussion

Nuclear localisation of AURKA has been implicated in its oncogenic activity in various cancers, where it has been associated with transcriptional regulation and chromatin remodelling^38–40^. These functions of AURKA are distinct from the well-characterised roles of AURKA kinase activity in mitosis^41–43^ and may be kinase-independent. Motivated by the possibility of using PROTACs for therapeutic manipulation of kinase-independent AURKA functions, and our previous observation of enhanced PROTAC sensitivity of a nuclear-localised mutant of AURKA^6^, we sought to investigate the sensitivity of nuclear AURKA to PROTAC activity. We found that forced nuclear localisation of AURKA (NLS-AURKA) enhanced its sensitivity to degradation by PROTACs, both by those binding to AURKA directly, and orthogonally using the dTAG system^35,36^. Since enhanced degradation of NLS-AURKA was not accompanied by any detectable increase in ternary complex formation between CRBN and NLS-AURKA (compared to WT-AURKA), we concluded that the increased degradation was most likely mediated through a post-ubiquitination step, such as the trimming or removal of ubiquitin chains by DUBs.

We therefore systematically investigated the effects of DUB knockdown on PROTAC-mediated degradation of AURKA. Previous studies have identified USP2 and OTUD6A as interactors and regulators of AURKA^29,30^ and both hits appeared in our DUBs-wide siRNA screen. Our subsequent validations confirmed OTUD6A as a key determinant of both cellular levels, and PROTAC sensitivity, of AURKA. Our screen also identified the proteasomal DUB UCHL5 as a limiting factor in PROTAC-mediated degradation. UCHL5 emerged as a broad regulator of PROTAC activity. Knockdown of UCHL5 enhanced the activity of all CRBN-recruiting PROTACs tested in our experiments (CRBN-recruiting AURKA-targeting PROTACs and dTAG-13) but had no effect on activity of VHL-recruiting dTAGv-1. UCHL5’s regulatory role may therefore depend on the type of ubiquitin chain topology induced by different E3s. These findings are consistent with the known proteasome-associated deubiquitinating activity of UCHL5^33,44^ acting to limit substrate processing for proteasomal degradation. While some previous studies have shown that UCHL5 knockdown can lead to enhanced degradation of substrates at the proteasome^45,46^, more recent studies have found instead that UCHL5 acts to promote substrate processing and degradation through removal of branched ubiquitin chains^47,48^. Our identification of UCHL5 as an E3-dependent PROTAC-opposing DUB aligns with the idea that its role in substrate processing is chain topology-dependent and acts to limit degradation of substrates ubiquitinated in response to PROTAC.

In contrast to UCHL5, OTUD6A exerted target-specific effects on degradation of AURKA by PROTACs. Knockdown of OTUD6A enhanced degradation of AURKA by PROTAC-D and TL12-186 but had no significant effect on degradation of HiBiT-FKBP12^F36V^ by dTAGs. Our data also reveals that OTUD6A regulates the sensitivity of AURKA to degradation by PROTACs in a manner dependent on AURKA subcellular localisation. OTUD6A localises exclusively to the cytoplasm, where it acts to limit the activity of PROTACs in bringing about degradation of AURKA. Nuclear-localised AURKA, in contrast to the cytoplasmic pool, is not protected from degradation and the nuclear pool is therefore more efficiently degraded in response to PROTAC treatments.

This differential regulation of the neosubstrate AURKA by OTUD6A underscores the importance of subcellular compartmentalisation in determining PROTAC efficacy. In the case of AURKA, our findings serve to rationalise therapeutic targeting of the nuclear oncogenic pool of AURKA via TPD.

More generally, our findings have broad implications for the development of PROTACs as therapeutic agents. Firstly, in incorporating a knowledge of subcellular localisation and DUB-specific interactions in the initial selection of targets suitable for a PROTAC approach. Secondly, in highlighting the potential to exploit DUB-specific interactions to finetune PROTAC activity at the subcellular level. Finally, in identifying DUB regulators of PROTAC efficacy, offering new opportunities for improving PROTAC performance through combination strategies, for instance transient inhibition of DUBs to enhance degradation of otherwise resistant targets. Through our investigation of the enhanced sensitivity of nuclear AURKA to PROTAC activity, we therefore provide new insight into how the activity of PROTACs can be limited in the cellular environment. Such studies will be critical for understanding the determinants of target specificity and for rationally designing PROTACs with improved therapeutic profiles.

## Materials and Methods

### Plasmids and molecular Cloning

pEGFP-OTUD6A, pEGFP-UCHL5 and pEGFP-UCHL5^C88A^ were the kind gifts of Prof Sylvie Urbé and Prof Stephen Jackson. The following sequences were obtained by gene synthesis (Genewiz from Azenta Life Sciences): siRNA-resistant OTUD6A, siRNA-resistant UCHL5, FKBP12^F36V^. siRNA-resistant OTUD6A and siRNA-resistant UCHL5 were inserted into the MCS of the pNeonN1 backbone. pNeonN1-AURKA-FKBP12^F36V^ and pNeonN1-FKBP12^F36V^ were created by inserting the FKBP12^F36V^ fragment into the pNeonN1-AURKA and pNeonN1 backbones respectively. Plasmids containing NLS were created by inserting the SV40 NLS fragment (PKKKRKV) into the appropriate backbone using restriction enzymes. The mNeon in all constructs is HA tagged at the C-terminus. pEGFP-OTUD6A^C152A^ was generated by round-the-horn site-directed mutagenesis of the pEGFP-OTUD6A plasmid using forward and reverse primers 5’-GCCATGTACCGCGCCATCCA-3’ and 5’-GTGGCCGTCGGCCGG-3’. NEB 5-alpha Competent E. coli (High Efficiency) (C2987I, NEB) were used. Full cloning details available upon request.

### Cell culture

U2OS parental and derived cell lines (U2OS FZR1^KO^, U2OS HiBiT-AURKA^TO^, U2OS^CDK2^) were cultured in DMEM (Thermo Fisher Scientific), supplemented with 10% V/V FBS (Sigma), 200 µM GlutaMAX-1 (Thermo Fisher Scientific), 100 U/ml penicillin, 100 µg/ml streptomycin (Gibco) and 250 ng/ml fungizone (Gibco). U2OS^CDK2^ were supplemented with 500 µg/ml G-418 disulfate salt solution (Sigma). U2OS HiBiT-AURKA^TO^ were supplemented with 150 µg/ml hygromycin (Fisher Scientific). Cells were grown at 37°C with 5% CO_2_. The inducible HiBiT-AURKA^TO^ cell line was generated from U2OS FRT/TO Flp-In^TM^ cells (Thermo Fisher Scientific) co-transfected with the pcDNA5 FRT/TO expression vector expressing the HiBiT-AURKA construct and pOG44 plasmid containing the FLP recombinase. 24-hours post-transfection cells were treated with 150 µg/ml hygromycin (Fisher Scientific) for 7 days then selected clones were picked and expanded. Expression of tetracycline-induced constructs was achieved by adding 1 µg/ml doxycycline (Sigma) to the media.

### Drug treatments

AURKA PROTACs were synthesised in-house at AstraZeneca^6^ and used at concentrations ≤ 10 µM. dTAG PROTACs (Bio-techne) were used at concentrations ≤ 1 µM. MG132 (Alfa Aesar) and CB-5083 (Stratech Scientific) were used at 5 µM.

### Cell transfection

For the siRNA screen, the ON-TARGET*plus* siRNA library for deubiquitinating enzymes (Horizon Discovery) was used, with SMARTpool siRNA comprised of 4 different oligos per DUB. U2OS HiBiT-AURKA^TO^ cells were reverse transfected with siRNA using FuGENE SI transfection reagent (Promega) under an optimised protocol. Analysis was carried out 48 hours post-transfection. For all other experiments, U2OS, U2OS FZR1^KO^ and U2OS^CDK2^ cells were electroporated using the mNeon Transfection System (Thermo Fisher Scientific) with parameters 1150V pulse voltage, 30 ms pulse width and two pulses. Analysis was carried out 24 hours post-transfection where plasmids only were transfected, or 48 hours post-transfection where siRNA was transfected.

siRNA oligos used: GL2i (non-targeting negative control) 5’-CGUACGCGGAAUACUUCGAUU-3’; OTUD6A siRNA #1 5’GCACUACAACUCCGUGACATT-3’; OTUD6A siRNA #2 5’-GAGAAAGAAUGGAGUCCGA-3’; UCHL5 siRNA #2 5’-GCAGUUAAUACCACUAGUA-3’.

### HiBiT lytic assay

U2OS HiBiT-AURKA^TO^ cells were seeded into white-walled 96- or 384-well plates at appropriate density. 16 hours prior to assay, expression of HiBiT-AURKA was induced with doxycycline. Following appropriate drug treatments of the cells, bioluminescence was read using the Nano-Glo® HiBiT Lytic Detection System (Promega). Briefly, cells were lysed in 100 µl of a pre-made lysis buffer containing LgBiT and NanoLuc substrate furimazine and the plate was left on a benchtop shaker for 15 minutes at 100 rpm at room temperature. Bioluminescence was then read on a PHERAstar or CLARIOstar Microplate Reader (BMG LABTECH). Unless specified otherwise, three biological repeats were carried out for each experiment and each experiment contained three technical replicates.

### Cell viability assay

Following appropriate drug treatments of cells, cell viability was measured using the CellTiter-Glo® assay (Promega). Briefly, media was removed from cells. 40 µL CellTiter-Glo® reagent was added to each well and the plate was left on a benchtop shaker for 15 minutes at 100 rpm at room temperature. Bioluminescence was then read on a PHERAstar Microplate Reader (BMG LABTECH).

### Immunoblot

Cell lysates were prepared in NuPage SDS sample buffer (Invitrogen) with 100 µM DTT. Samples were boiled and loaded with Molecular weight PageRuler Prestained Protein Ladder into SDS-PAGE 4-12% Bis-Tris pre-cast gels (ThermoFisher) and run in 1XNuPAGE MOPS SDS Running Buffer (Invitrogen) (80 minutes, 150V). A semi-dry Pierce^TM^ G2 Fast Blotter was used for transfer to a Immobilon-FL PVDF membrane (Sigma) at 25V, 1.3A for 12 minutes according to the manufacturer’s instruction. Blocking and primary antibody incubations were performed in phosphate-buffered saline (PBS), 0.1% Tween-20, 5% low-fat milk overnight at 4°C. Signals were quantified by enhanced chemiluminescence detection, or using fluorophore-conjugated secondary antibodies, scanned on an Odyssey® Imaging System (LI-COR Biosciences).

Primary antibodies for immunoblot were as followed: AURKB rabbit polyclonal Ab (1:1000, Abcam 2254), AURKA mouse monoclonal Ab (1:1000, BD Biosciences 610938), Vinculin mouse monoclonal Ab (1:1000, Sigma 9131),OTUD6A rabbit polyclonal Ab (1:1000, Proteintech), UCHL5 rabbit monoclonal Ab (1:1000, Abcam ab133508), GAPDH mouse monoclonal Ab (1:4000, Proteintech 6004-1-Ig). HA rabbit polyclonal Ab (1:1000, Cell Signalling Technology 3724), HA mouse monoclonal Ab (1:1000, Cell Signalling Technology 2367).

Secondary antibodies used: Polyclonal Goat Anti-Rabbit or Polyclonal Rabbit Anti-Mouse (1:5000) HRP-conjugated (Dako Agilent), or IRDye® 680RD (1:10,000)- or 800RD (1:10,000)-conjugated for quantitative fluorescence measurements on an Odyssey® Fc Dual-Mode Imaging System (LICOR Biosciences). IRDye® conjugated antibodies were prepared in PBS, 0.1% Tween-20, 5% FBS, 0.01% SDS.

### Ubiquitin pulldown assays

Cells were electroporated with pcDNA3-FLAG-Ub, plasmids expressing mNeon-HA-tagged versions of AURKA and different siRNAs. Cells were treated with appropriate drugs for 1 hour before harvesting by scraping into RIPA lysis buffer containing protease and phosphatase inhibitors (ThermoFisher) and 50 mM NEM. Lysates were sonicated and incubated on ice for 1 hour, followed by centrifugation and collection of supernatant. 1 mg of sample was mixed with 20 µl Ubiquitin Affinity Beads (Universal Biologics Ltd.) at 4°C for 2 hours with rotation. Beads were collected by centrifuging at 5,000 rpm for 1 minute at 4°C. Beads were washed 3 times with 100 µL ice-cold wash buffer (50% lysis buffer). Protein was eluted in non-reducing sample buffer with a 3-minute incubation at 95°C. Beads were centrifuged at 13,000 rpm at 4°C for 1 minute and lysate was transferred to a clean Eppendorf containing 1 µL β-mercaptoethanol. Eluate and input samples were boiled at 95°C for 5 minutes prior to SDS-PAGE gel electrophoresis and immunoblotting for detection of ubiquitin-conjugated substrate with HA antibody.

### Immunofluorescence

Cells were seeded onto glass coverslips in 6-well tissue culture plates at a density of 250,000 cells per well. 24 hours later, cells were fixed by addition of -20 °C methanol. On the day of staining, cover slips were permeabilized, blocked in blocking buffer (0.1% (v/v) Triton-X100, 3% (w/v) BSA, PBS) for 15 minutes at room temperature, washed three times in PBS, 0.1% Triton and transferred to a humidity chamber. Primary antibodies were diluted in blocking buffer and added to coverslips for a 90-minute incubation at room temperature. Cover slips were washed three times in PBS, 0.1% Triton, followed by a one-hour incubation with secondary antibody, again diluted in blocking buffer. Cells were washed once in PBS, 0.1% Triton, then once in PBS containing 1 µg/ml DAPI, then once in PBS. Cover slips were mounted onto microscope slides using ProLong Gold anti-fade mounting media (ThermoFisher).

Primary antibodies used: AURKA mouse monoclonal Ab (1:1000, clone 4/IAK1, BD Transduction Laboratories), beta-tubulin rabbit monoclonal Ab (1:1000, Abcam ab6046). Secondary antibodies used: Alexa Fluor 488 anti-mouse and Alexa Fluor 568 anti-rabbit (Thermo Fisher Scientific).

### *is*PLA

Cells were seeded onto coverslips, fixed with paraformaldehyde and *is*PLA was carried out using the Duolink PLA kit (DUO92004 and DUO92002; Sigma-Aldrich) according to manufacturer’s instructions, with an amplification time of 80 minutes. The primary antibody pair to detect the interaction was mouse anti-Neon (1:1000, Proteintech 32F6)/rabbit anti-CRBN (1:500 Abcam ab226782), and DNA was stained with DAPI.

### Microscopy

Live-cell microscopy was carried out in L-15 medium (Life Technologies) supplemented with 10% V/V FBS on an Olympus CellR platform made up of an Olympus IX81 motorized inverted microscope with a 40X NA 1.3 oil immersion objective lens, a Retiga6-CCD Camera (Photometrics), a motorised stage (Prior Scientific), CoolLED pE-4000 illumination system and a 37°C incubation chamber (Solent Scientific). Images were acquired using appropriate LED lines and filter sets under control of Micromanager software^49^ and exported as 16-bit tiff files. Quantitative image analysis was achieved using FIJI software^50^. The total fluorescence of single cells was calculated from ROIs drawn around each cell, measuring total fluorescence corrected for background measured outside the cell. For calculation of the nuclear to cytoplasmic ratio of the mNeon signal for WT- and NLS-AURKA-Neon constructs and of the Cerulean signal in U2OS^CDK2^ cells, a customised plug-in tool (Multi_Measure) was used to calculate mean fluorescence intensity (MFI) by measuring average, background-subtracted grey values of ROIs of defined diameter (15 µm) around manually selected points in the cell. Immunofluorescence slides were imaged using an Olympus IX83 motorized inverted microscope with a 40X air lens, Spectra-X multi-channel LED widefield illuminator (Lumencor), Optospin filter wheel (Cairn Research), CoolSnap MYO CCD camera (Photometrics), automated XY stage (ASI) and climate chamber (Digital Pixel). Images were acquired as stacked TIFF format using MicroManager software. The TIFF files were imported into FIJI and the z-stacks were projected based on maximum intensity. Total fluorescence of single cells was calculated from ROIs drawn around each cell, subtracting background fluorescence measured outside of the cell.

### Statistical analysis

GraphPad Prism 10 (version 10.0.3; GraphPad Software Inc) and Microsoft Excel (version 16.78; Microsoft Corporation) were used to analyse data, generate graphs and perform statistical analysis. For datasets with two groups, t-test was used to assess significance. For datasets with multiple groups, one-way ANOVA was used. Two-way ANOVA was used to assess significance in datasets with two factors. The number of biological repeats and statistical parameters are indicated in figure legends. For all statistical tests applied, results are indicated as p < 0.05 (*), p ≤ 0.01 (**), p ≤ 0.001 (***), p ≤ 0.0001 (****).

## Additional Information

### Author Contributions

A.C. Conceptualisation, Data curation, Formal analysis, Investigation, Methodology, Writing – original draft, Writing – review and editing; K.R. Conceptualisation, Writing – review and editing; C.L. Conceptualisation, Funding acquisition, Writing – original draft, Writing – review and editing.

## Acknowledgements

We thank past and present members of the Lindon lab for insightful discussions of the study. We extend thanks to Linda Kitching and Eleni Charla at AstraZeneca for their guidance in setting up the screening experiments. We are grateful to Prof Sylvie Urbé and Prof Stephen Jackson for providing essential plasmids. Special thanks to Amélie Davies and Karthik Sadanand for their contributions to the cell cycle analysis of DUB knockdowns. Schematic illustrations were created in BioRender.

---

**Supplementary Figure 1:**
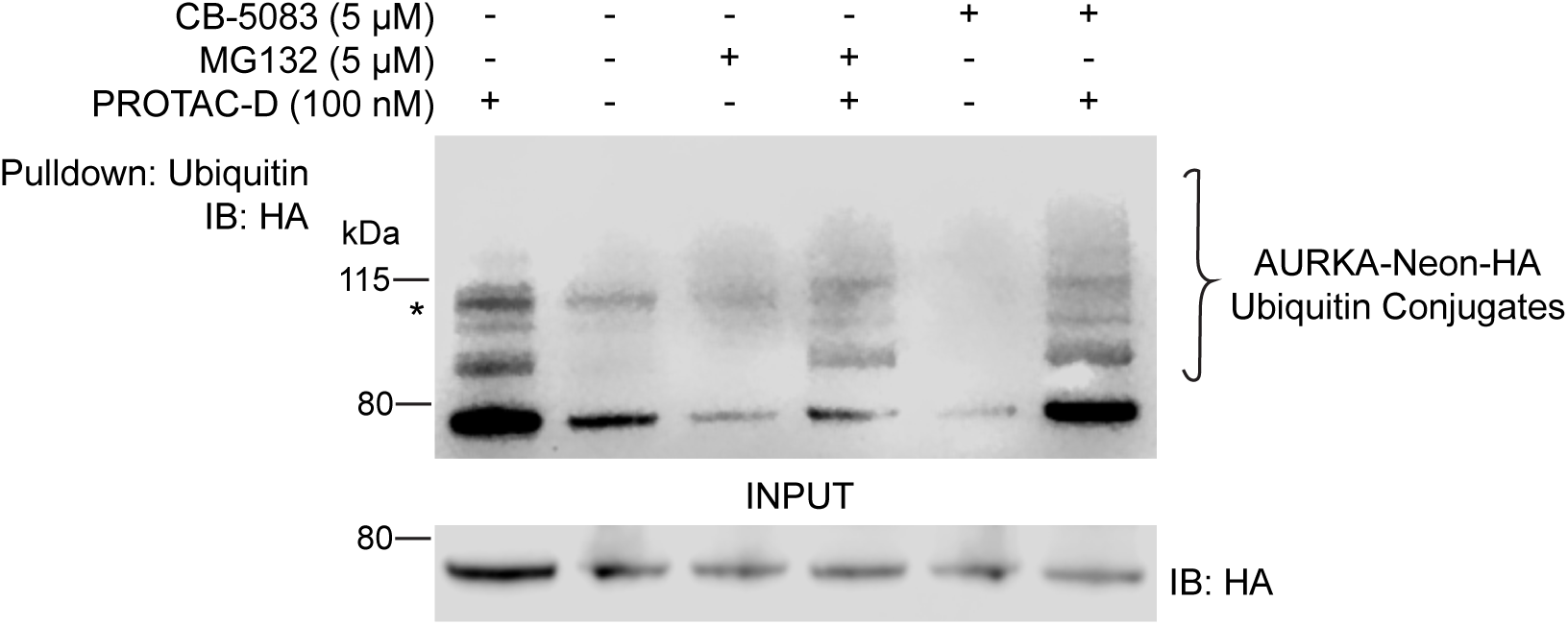
PROTAC-induced ubiquitination of AURKA-Neon-HA. U2OS FZR1^KO^ cells co-transfected with pNeonN1-AURKA-HA and pcDNA3 FLAG-Ub were treated as indicated for one hour. Ubiquitinated proteins from cell extracts were captured by pulldown with ubiquitin affinity beads. The band marked (*) is a non-specific band. Treatment with p97 inhibitor (CB-5083), but not proteasome inhibitor (MG132), enriched pulldowns for polyubiquitinated AURKA.

**Supplementary Figure 2:**
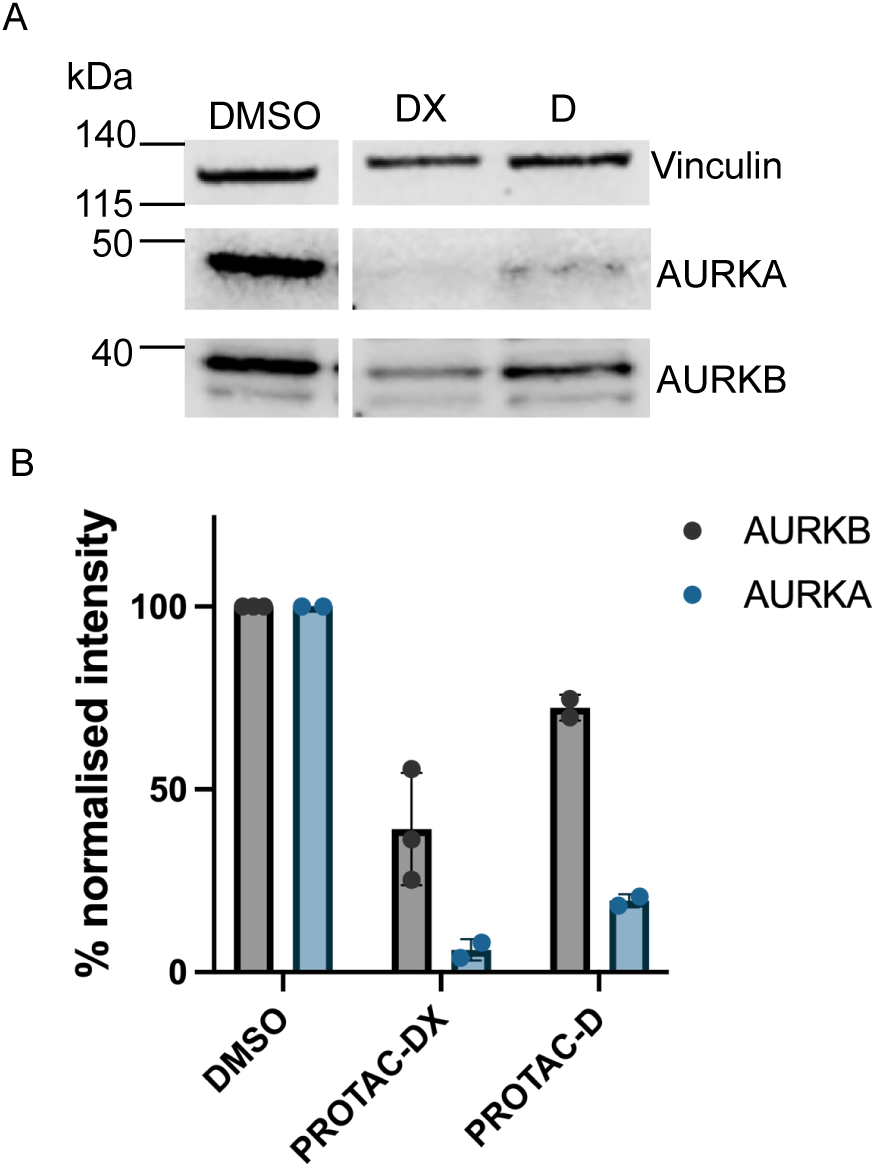
AURKB is sensitive to PROTAC-DX. **A** Immunoblot of U2OS cells treated with DMSO, PROTAC-DX or PROTAC-D (1 µM) for 4 hours prior to lysate collection. Panels are from the same immunoblot. **B** Quantification of vinculin-normalised AURKA and AURKB levels from the immunoblot in (**A**). Bar chart shows mean values ± SD from two or three biological repeats.

**Supplementary Figure 3:**
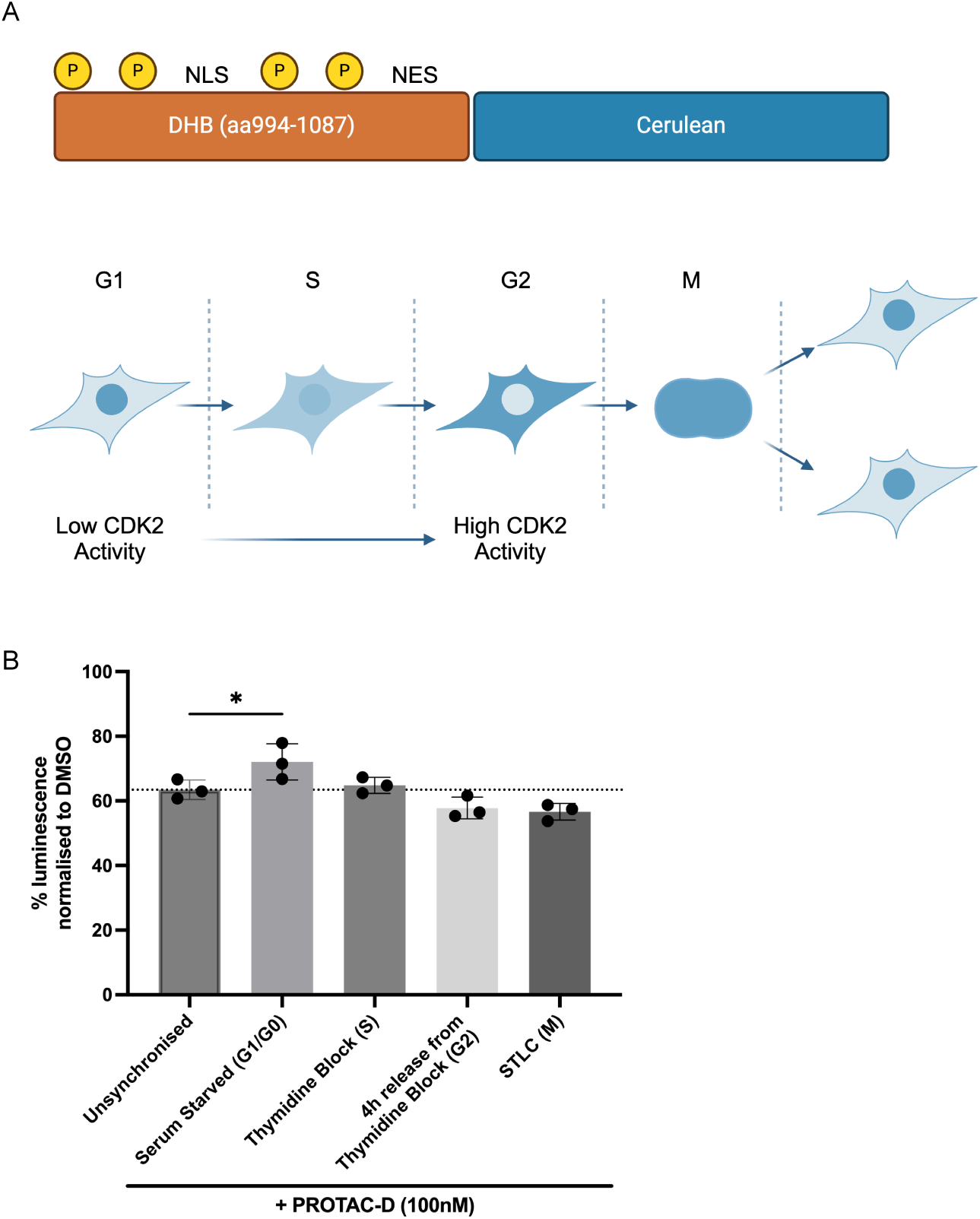
Cell cycle influence on PROTAC-D activity. **A** Schematic representation of the CDK2 activity sensor expressed in U2OS^CDK2^ cells. **B** U2OS HiBiT-AURKA^TO^ cells were synchronised as indicated and treated for 4 hours with DMSO or PROTAC-D (100 nM). Bar chart shows DMSO-normalised HiBiT-AURKA levels as mean values ± SD from n=3 biological repeats; statistical analysis by one-way ANOVA with Dunnett’s post hoc multiple comparisons to unsynchronised cells.

**Supplementary Figure 4:**
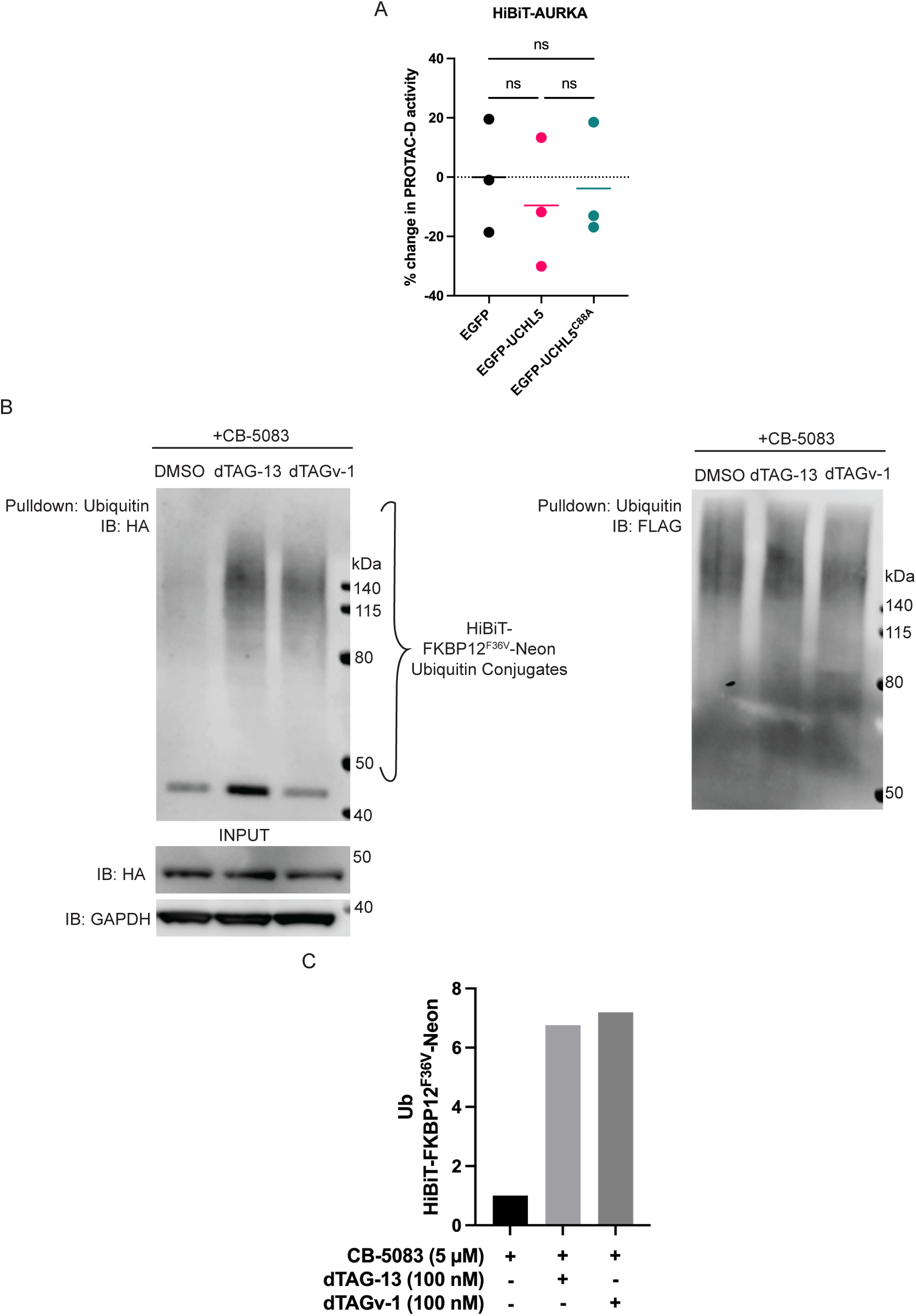
PROTAC-induced ubiquitination of HiBiT-FKBP12^F36V^-Neon-HA. **A** U2OS HiBiT-AURKA^TO^ cells transfected with pEGFP, pEGFP-UCHL5, or catalytically inactive pEGFP-UCHL5^C88A^ plasmids were treated with DMSO or PROTAC-D (100 nM) for 4 hours. Data points show percentage change in PROTAC-mediated HiBiT-AURKA degradation relative to pEGFP overexpression and correspond to mean values of technical replicates from n=3 biological repeats with line to indicate the mean value of biological repeats; statistical analysis by one-way ANOVA with Tukey’s post hoc multiple comparisons test. **B** U2OS cells co-transfected with pNeonN1-HiBiT-FKBP12^F36V^ and pcDNA3-FLAG-Ub were treated with DMSO, dTAG-13 or dTAGv-1 (100 nM) and CB-5083 (5 µM) for one hour. Ubiquitinated proteins from cell extracts were captured by pulldown with ubiquitin affinity beads. **C** Quantification of (**B**): chart shows the ratio of ubiquitin conjugates to input for each condition, normalised to the ratio for DMSO treatment.

